# UK phenomics platform for developing and validating EHR phenotypes: CALIBER

**DOI:** 10.1101/539403

**Authors:** Spiros Denaxas, Arturo Gonzalez-Izquierdo, Kenan Direk, Natalie K Fitzpatrick, Ghazaleh Fatemifar, Amitava Banerjee, Richard Dobson, Laurence J. Howe, Valerie Kuan, R. Tom Lumbers, Laura Pasea, Riyaz S. Patel, Anoop D Shah, Aroon D. Hingorani, Cathie Sudlow, Harry Hemingway

## Abstract

**Objective:** Electronic Health Records (EHR) are a rich source of information on human diseases, but the information is variably structured, fragmented, curated using different coding systems and collected for purposes other than medical research. We describe an approach for developing, validating and sharing reproducible phenotypes from national structured EHR in the United Kingdom (UK) with applications for translational research.

**Materials and Methods:** We implemented a rule-based phenotyping framework, with up to six approaches of validation. We applied our framework to a sample of 15 million individuals in a national EHR data source (population-based primary care, all ages) linked to hospitalization and death records in England. Data comprised continuous measurements e.g. blood pressure, medication information and coded diagnoses, symptoms, procedures and referrals, recorded using five controlled clinical terminologies: a) Read (primary care, subset of SNOMED-CT), b) International Classification of Diseases 9th/10th Revision (ICD-9, ICD-10, secondary care diagnoses and cause of mortality), c) OPCS Classification of Interventions and Procedures (OPCS-4, hospital surgical procedures), and d) DM+D prescription codes.

**Results:** Using the CALIBER phenotyping framework, we created algorithms for 51 diseases, syndromes, biomarkers and lifestyle risk factors and provide up to six validation approaches. The EHR phenotypes are curated in the open-access CALIBER Portal (https://www.caliberresearch.org/portal) and have been used by 40 national/international research groups in 60 peer-reviewed publications.

**Conclusion:** We describe a UK EHR phenomics approach within the CALIBER EHR data platform with initial evidence of validity and use, as an important step towards international use of UK EHR data for health research.

## BACKGROUND AND SIGNIFICANCE

The United Kingdom (UK) National Health Service (NHS) offers international researchers opportunities to explore ‘cradle to grave’ longitudinal electronic health record (EHR) phenotypes at scale. It is one of the few countries which combines a single-payer-and-provider comprehensive healthcare system, free at the point of care, with extensive national data resources across the entire 65M population. Patients are identified by a unique healthcare-specific identifier which enables linkage of patient data across EHR sources and the creation of longitudinal phenotypes that span primary and secondary care [1]. Over 99% of people are registered with a general practitioner (GP) and structured primary care data collected electronically have been used by UK, United States (US) and other researchers for decades [2]. Furthermore, these national EHR data sources are being linked with large-scale consented genomic resources i.e. 100,000 Genomes Project (also known as Genomics England) [3] and UK Biobank [4–6] and enable the investigation of simple/complex traits across participant populations with diverse genetic backgrounds [7].

The UK EHR landscape differs from the US and elsewhere in important ways. Although the UK, unlike the US, has the opportunity to establish a national approach, it faces the common challenge that EHR for primary care and hospital care are handled by different data providers and are kept separately, with independent access requirements [8,9]. Significant progress has been made by US initiatives (Electronic Medical Records and Genomics (eMERGE) [10], BioVU [11], Million Veteran Programme (MVP), [12] *All Of Us* [13]), Canada [14], Australia [15], Sweden [16] and Denmark [17]. In the UK however, there has been no recognized phenotyping framework or go-to resource for EHR researchers for systematically creating, curating and validating (rule-based or otherwise) EHR-derived phenotypes, obtaining information on controlled clinical terminologies, sharing algorithms, and communicating best approaches. Structured primary care EHR have been used in >1800 published studies [18] but only 5% of studies published sufficiently reproducible phenotypes [19] while significant heterogeneity exists (one review reported 66 asthma definitions [20]). Current UK initiatives [19,21,22] for curating EHR-derived phenotypes focus on lists of controlled clinical terminology terms (referred to as *codelists*) rather than self-contained phenotypes i.e. terms, implementation, validation evidence.

The scope of our research focuses on rule-based algorithms as the majority of research studies (with some exceptions [23,24]) using UK EHR utilize this approach for creating EHR-derived phenotypes [25]. The main use-case for CALIBER phenotypes and the approach presented in the manuscript is observational research (which is also the main stakeholder group of UK EHR): a) high-resolution clinical epidemiology using national EHR examining disease aetiology or prognosis, or b) genetic epidemiology studies through the UK Biobank and Genomics England investigating simple and complex traits across populations. Our aspiration however is for CALIBER phenotypes to be adopted by the NHS in terms of computable knowledge which can be integrated in the healthcare system and used for interventional studies and clinical guidelines. Each of these use cases however has a different threshold on what is considered adequate performance and we adopted a systematic and robust validation approach in order to quantify phenotype performance.

EHR phenotype validation is a critical process guiding their subsequent use in research or care [26,27]. There are multiple sources of evidence/study designs that contribute to building confidence in the validity of an EHR phenotype for a particular purpose. Countries may also differ in the opportunities for validation: e.g. in the UK cross-referencing against multiple EHR sources, prognostic validation and risk factor validation are all made possible by nationwide population-based records [28–32]. In contrast with the US, only recently have scalable methods been developed to access the entire hospital record for expert review [33] and text corpora are not available at scale [34]. There have been few previous studies [35] of the validity of International Classification of Disease and Health Related Problems, 10th Revision (ICD-10) terms [36] in the UK against hospital records because introduction of hospital EHRs are recent (e.g. there are only three hospitals that have achieved stage six on the Healthcare Information and Management Systems Society (HIMSS) Electronic Medical Record Adoption Model (EMRAM) [37]).

We have developed the CALIBER EHR platform for the UK by adopting and extending best practices from leading initiatives and consortia (e.g. eMERGE, MVP, BioVU and others) with regards to creating, evaluating and disseminating EHR-derived phenotypes for research. Specifically, these practices, which were previously not systematically followed in the UK EHR community prior to CALIBER, include: a) establishing a robust and iterative phenotype creation process involving multiple scientific disciplines, b) systematically curating EHR-derived phenotypes, c) using methods for enhancing reproducibility, and d) undertaking and reporting robust phenotype validation analyses. Here, we define a framework for enabling EHR phenotyping in a scalable and reproducible manner. Algorithm reproducibility was defined similarly to Goodman’s “methodology reproducibility” [38] i.e. providing a systematic and precise description of the algorithm components, logic, implementation and evidence of validity that would enable national or international independent researchers to create, apply and evaluate CALIBER phenotyping algorithms in local similar data sources. We present a systematic validation framework for assessing accuracy consisting of up to six approaches of evidence (i.e. expert review to prognostic validation) and disseminating through a centralized open-access repository. We have chosen heart failure (HF), acute myocardial infarction (AMI) and bleeding as examples of medical conditions that exemplify the strengths of national linked UK EHR and the non-trivial challenges researchers encounter.

## MATERIALS AND METHODS

We developed an iterative and collaborative approach for creating and validating rule-based EHR phenotyping algorithms using UK structured EHR. The approach involved expert review interwoven with data exploration and analysis. An EHR phenotyping algorithm translates the clinical requirements for a particular patient to be considered a case into queries that leverage EHR sources stored in a relational database and extracts disease onset, severity and subtype information. In the following sections we describe the platform, the algorithm development process and validation consisting of six approaches of evidence.

### UK primary care EHR, hospital billing data and cause-specific mortality in the CALIBER platform

The CALIBER platform [39] is currently built around four national EHR data sources **(Figure 1)** deterministically linked using NHS number (unique ten-digit identifier assigned at birth or first interaction), gender, postcode and date of birth; 96% of patients with a valid NHS number successfully linked [40].

The baseline cohort is composed of a national primary care EHR database, the *Clinical Practice Research Datalink (CPRD) [41]*. Primary care has used computerised health records since 2000 and general practices use one of several EHR systems. CPRD contains longitudinal primary care data (extracted from the Vision and Egton Medical Information Systems (EMIS) clinical information systems) on diagnoses, symptoms, drug prescriptions, vaccinations, blood tests and risk factors (irrespective of disease status and hospitalization). CPRD uses Read [42] terms (112,806 terms, subset of the The International Health Terminology Standards Development Organisation Systematized Nomenclature Of Medicine-Clinical Terms (SNOMED-CT) [42]) to record information. Prescriptions are recorded using Gemscript (a commercial derivative of the NHS Dictionary of Medicines and Devices (DM+D)) [43] (72,664 entries). CPRD contains >10billion rows of data from >15M patients (from all the contributing primary care practices, irrespective of consent to linkage) shown to be representative in terms of age, sex, mortality and ethnicity [44–46] and of high validity [47].

*Hospital Episode Statistics (HES)* (https://digital.nhs.uk/) [48] contains administrative data on diagnoses and procedures generated during hospital interactions. Diagnoses are recorded using the International Classification of Diseases 10th Revision (ICD-10) and procedures using The Office of Population Censuses and Surveys Classification of Surgical Operations and Procedures, 4th Revision (OPCS-4) (10,713 terms, similar to Current Procedural Terminology (CPT) [49]). Up to 20 primary and secondary discharge diagnoses are recorded per finished consultant episode. The *Myocardial Ischaemia National Audit Project (MINAP)*, is a national disease and quality improvement registry capturing all acute coronary syndrome events across England. MINAP contains diagnostic, severity and treatment information using 120 structured data fields [50]. The *Office for National Statistics* (ONS) contains socioeconomic deprivation using the Index of Multiple Deprivation (IMD) [51] and physician-certified cause-specific mortality (underlying and up to 14 secondary causes using ICD-9/ICD-10).

### Data quality

#### Primary care

Our analyses incorporated primary care EHR data quality metrics across two dimensions: at the patient level and at the primary care practice level [41]:

##### Patient-level data quality

In line with previous research using UK primary care electronic health records from the Clinical Practice Research Datalink (CPRD) and CPRD guidance, we only utilized patients which were marked as “acceptable for research” by the CPRD. Patients are labelled as acceptable through an algorithmic process which identified and excludes patients with non-continuous follow up and patients with poor data according to a predefined list of data quality metrics (e.g. empty date of first registration, first registration prior to date of birth, invalid gender, missing or incorrect dates across all recorded healthcare episodes). We additionally excluded records where the date was invalid/malformed or in the future occurring after the last date of data collection.

##### Practice-level

The overall quality of the data recorded in a primary care practice is algorithmically marked by an “up to standard (UTS)” date by the CPRD. The UTS date is deemed as the date at which data in the practice is considered to have continuous high-quality data fit for use in research. The algorithm used to derive this date is based on two concepts: a) gap analysis (assurance of continuity in data recording and establishing if any unexpected and prolonged gaps in recording exist) and, b) death recording (observing the expected and actual deaths recorded at a practice over time by taking into account season and geographical variation in death rates and establishing if any gaps in recording exist). In both of these cases, the UTS date is set to the latest of these dates.

Completeness patterns of key clinical covariates such as risk factors (e.g. smoking status, blood pressure, BMI) has been previously shown to have rapidly increased after the introduction of a financial incentives framework (Quality and Outcomes Framework) which encourages GPs to record key data items [41].

#### Secondary care

Hospital Episode Statistics (HES) Admitted Patient Care (APC) data are collected for all admissions to all National Health Service (NHS) secondary healthcare providers. The NHS funds 98-99% of hospital activity in England. HES APC are administrative data collected for reimbursement of hospital activity and are post-discharge derived by clinical coders according to standardized rules for translating information from from discharge summaries into diagnosis (ICD-10) and surgical procedure terms (OPCS-4) terms [48]. The overarching reimbursement framework, Payment-By-Results (a fixed tariff case mix based payment system [52]), provides financial incentives for hospitals to improve their coding accuracy and depth and ensure accurate reimbursement. This has led to an increase in the number of diagnosis terms recorded and coding accuracy i.e. primary diagnoses accuracy was 96% (interquartile range (IQR): 89.3-96.3) when compared to expert review of case notes [53]. The NHS Digital Data Quality Maturity Index (DQMI) provides a per hospital overall score for clinical data quality in term of data field and hospitalization episode completeness on a quarterly basis [54].

### Algorithm development

The development pipeline was a collaborative and iterative process involving researchers from a diverse set of scientific backgrounds (e.g. clinicians, epidemiologists, computer scientists, public health researchers, statisticians). An iteration refers to an adjustment in the computational strategy to derive the phenotype in question, based on data-driven examinations of its internal validity and according to the clinical context. The number of development iterations was proportionate with the complexity of the clinical phenotype: algorithms leveraging multiple sources required multiple iterations and substantially more clinician input.

We initially defined search strategies for identifying relevant diagnosis terms and their synonyms which were selected based on input from clinicians, existing literature, national guidelines and by consulting medical vocabulary repositories e.g. Unified Medical Language System (UMLS) Metathesaurus [55,56]. Two clinicians independently classified identified terminology terms (disagreements resolved by third) into non-overlapping categories: a) *prevalent* (e.g. “history of heart failure”) b) *possible* (e.g. “congestive heart failure monitoring”), and c) *incident* (potentially sub-classified e.g. “chronic congestive heart failure”, “acute left ventricular failure”, “heart failure not otherwise specified”). Similarly, we identified and classified coded symptoms recorded in primary care EHR. Many CALIBER phenotyping algorithms combine coded diagnosis, symptom information, continuous measurements e.g. laboratory values or other physiological measurements and medication prescription information in a rule-based fashion e.g. hypertension is defined using continuous blood pressure, coded diagnoses, blood-pressure lowering prescriptions, and comorbidities. We generated ad-hoc rules to reconcile: a) coding differences across EHR sources with respect to the granularity of diagnosis, b) the presence of multiple terms i.e. multiple different ethnicity entries, and c) transience in coding (e.g. ICD-9 was used for recording the cause of death before 2000). In primary care EHR, identified Read terms were evaluated in terms of their information content and subsequent ability to ascertain a phenotype reliably.

Primary care EHR contain over 450 structured data items for recording continuous measurements e.g. blood markers. For continuous phenotypes (e.g. blood pressure), we normalised data quality by identifying the relevant units, specified unit conversions (where required) and defined valid value ranges. For example, the neutrophil count structured data area contained both the absolute values and percentages, and these had to be differentiated by supplementary Read terms and by checking the distribution of values by unit. Sometimes values were obviously on the wrong scale e.g. haemoglobin where some values were distributed as if measured in g/L but had (presumably incorrect) units recorded as g/dL. Zero values caused particular problems; they could be impossible and represent missing data in some cases (e.g. ferritin) but might be true zeroes representing undetectable values in other cases (e.g. basophils). Careful investigation by units and Read term was required to avoid creating Missing Not at Random data (if the zeroes were true) or false data (if the zeroes were false). Definition of valid ranges for values was also problematic, as we wanted to exclude erroneous values without excluding true physiologically extreme values.

### Validation: Systematic evaluation using six approaches

Obtaining and curating evidence of phenotype validity is an essential component of the phenotyping process. We evaluated EHR-derived phenotypes across up to six different approaches of providing of evidence of phenotype validity, acknowledging that that the use case will inform which validation(s) are most important. For example, phenotyping algorithms developed for disease epidemiology (e.g. screening or disease surveillance) might be designed for higher sensitivity whereas those used in genetic association studies might be designed to maximize PPV. We provide details of these validation approaches below:

#### 1) Cross-EHR source concordance

For EHR-derived cases of AMI, HF and bleeding, we quantified the percentage of cases identified in each source, the overlap between sources and evaluated per-source completeness and diagnostic validity. Additionally, we used a disease registry (MINAP) as a reference in order to derive the positive predictive value (PPV) of AMI diagnoses recorded in hospital EHR (HES) i.e. the probability that an AMI diagnosis recorded in HES was indeed an AMI as ascertained by MINAP (that contains information on AMI ascertainment such as electrocardiogram results and troponin measurements) rather than unstable angina or a non-cardiac diagnosis. We did not calculate sensitivity and specificity relative to MINAP given that MINAP does not include all cases of AMI, as it is a disease registry which requires bespoke data entry by audit staff separate from clinical care or coding. It is therefore not possible to use MINAP as a gold standard to evaluate hospital EHR (Hospital Episode Statistics) in relation to completeness of detection of AMI (sensitivity) or non-MI (specificity). However, there is a concern that HES data may be inaccurate, and MINAP can be used to evaluate its positive predictive value for the subset of cases with a MINAP record for the event, where the exact diagnosis in MINAP can be considered a “gold standard.”

#### 2) Case note review

We evaluated the performance of the secondary care component of the bleeding phenotype by assessing the ability of the diagnosis terms (ICD-10) utilized by the phenotype to correctly identify hospitalized bleeding events in two independent hospital EHR sources. Two clinicians (blinded to the ICD-10 diagnosis terms) reviewed the entire hospital record (charts, referral letters, discharge letters, imaging reports) for 283 completed patient hospital episodes across two large hospitals (University College London Hospitals, King's College Hospital). Bleeding assignments from the clinicians review was compared with those from the phenotyping algorithm and we estimated the PPV, NPV, sensitivity and specificity using the case review data as the “gold standard”. We extracted hospital data (14,364,947 words) using CogStack [57] from the consented Stroke InvestiGation Network-Understanding Mechanisms (SIGNUM) study.

#### 3) Consistency of risk factor-disease associations from non-EHR studies

For all exemplars, we produced and reported hazard ratios (HR) and 95% Confidence Intervals (CI) of known risk factors from Cox proportional hazards models adjusted for age, sex and other covariates). We evaluated the ability of obtaining consistent estimates (in terms of direction and magnitude) with risk factor associations derived from non-EHR research-driven studies.

#### 4) Consistency with prior prognosis research

We produced Kaplan–Meier (KM) cumulative incidence curves at appropriate time intervals and endpoints and stratified by EHR source. We evaluated the observed prognostic profiles with previously-reported evidence for example observing different survival patterns between patients diagnosed with HF in CPRD but never hospitalized compared with patients diagnosed in HES.

#### 5) Consistency of genetic associations

Similar to previous studies, we attempted to replicate previously reported associations between genetic variants and diseases discovered from non-EHR studies (e.g. research-driven observational cohort studies or interventional studies). The ability of EHR-derived phenotypes to replicate previously-discovered associations derived from non-EHR studies and observing similar direction and magnitude of association reinforces the evidence towards the overall validity of the EHR phenotype [58]. Using PLINK [59], we extracted genetic variants associated with AMI reaching genome-wide significance (P <5×10^−8^) from publicly-available 1000 Genomes-based Genome Wide Association Study (GWAS) summary data (“CARDIoGRAMplusC4D - mi.additive.Oct2015”) in the CARDIoGRAMplusC4D [60] consortium. In the UK Biobank, we identified AMI cases in linked hospital and mortality EHR using the CALIBER AMI phenotype and defined controls as non-case participants with no self-reported record of AMI at baseline. We estimated the association of genetic variants and AMI using logistic regression with an underlying additive model in PLINK adjusting for the first 10 principal components, age and sex. Replication was defined as the Single Nucleotide Polymorphism (SNP) being associated with AMI in the UK Biobank (Bonferroni-adjusted P<0.0016) with a concordant direction of effect with CARDIOGRAMPlusC4D.

#### 6) External populations

We assessed the validity of developed algorithms by implementing them in external data sources (UK or elsewhere), and examining consistency of results in the evaluation criteria.

### Phenotype dissemination

We generated textual descriptions of algorithms with explicit detail on the logic behind the algorithm (pre-processing, cross-source reconciliation, quality checks) in a clinician-friendly manner. We generated flow-chart representations accompanied by pseudo-code for facilitating the translation of the algorithm to Structured Query Language (SQL) queries. We created entries in the CALIBER Portal **(Figure 4)** describing implementation details across sources, research outputs, validation evidence and a Digital Object Identifier (DOI) [61]. We created an open-source R library for manipulating clinical terminologies (http://caliberanalysis.r-forge.r-project.org/) using a custom file format including metadata (e.g. naming, version, authors, timestamp).

### Ethical approval

The CPRD has broad ethical approval for purely observational research using pseudonymised linked primary/secondary care data for supporting medical purposes that are in the interests of patients and the wider public. Linkages were performed by NHS Digital, the statutory body in England responsible for providing core healthcare information technology and curating many of the national datasets. This study was approved by the Medicines and Healthcare Products Regulatory Agency (MHRA) Independent Scientific Advisory Committee (ISAC) - protocol references: 11_088, 12_153R, 16_221, 18_029R2, 18_159R.

## RESULTS

Using the CALIBER EHR phenotyping approach described here, we curated over 90,000 terms from five controlled clinical terminologies to create 51 validated phenotyping algorithms (35 diseases/syndromes, ten biomarkers, six lifestyle risk factors). In this manuscript, we used three exemplar phenotypes: heart failure (https://www.caliberresearch.org/portal/phenotypes/heartfailure), bleeding (https://www.caliberresearch.org/portal/phenotypes/bleeding) and acute myocardial infarction (https://www.caliberresearch.org/portal/phenotypes/acutemyocardialinfarction). Table 1 provides a complete list of published, peer-reviewed phenotypes and the approaches of evidence supporting their validity. CALIBER phenotypes have been used by 40 national/international research groups in 60 peer-reviewed publications [62]. The CALIBER Portal (http://www.caliberresearch.org/portal) opened in October/2018 to the community and provides a centralized resource for curating EHR-derived phenotypes.

**Table 1:**
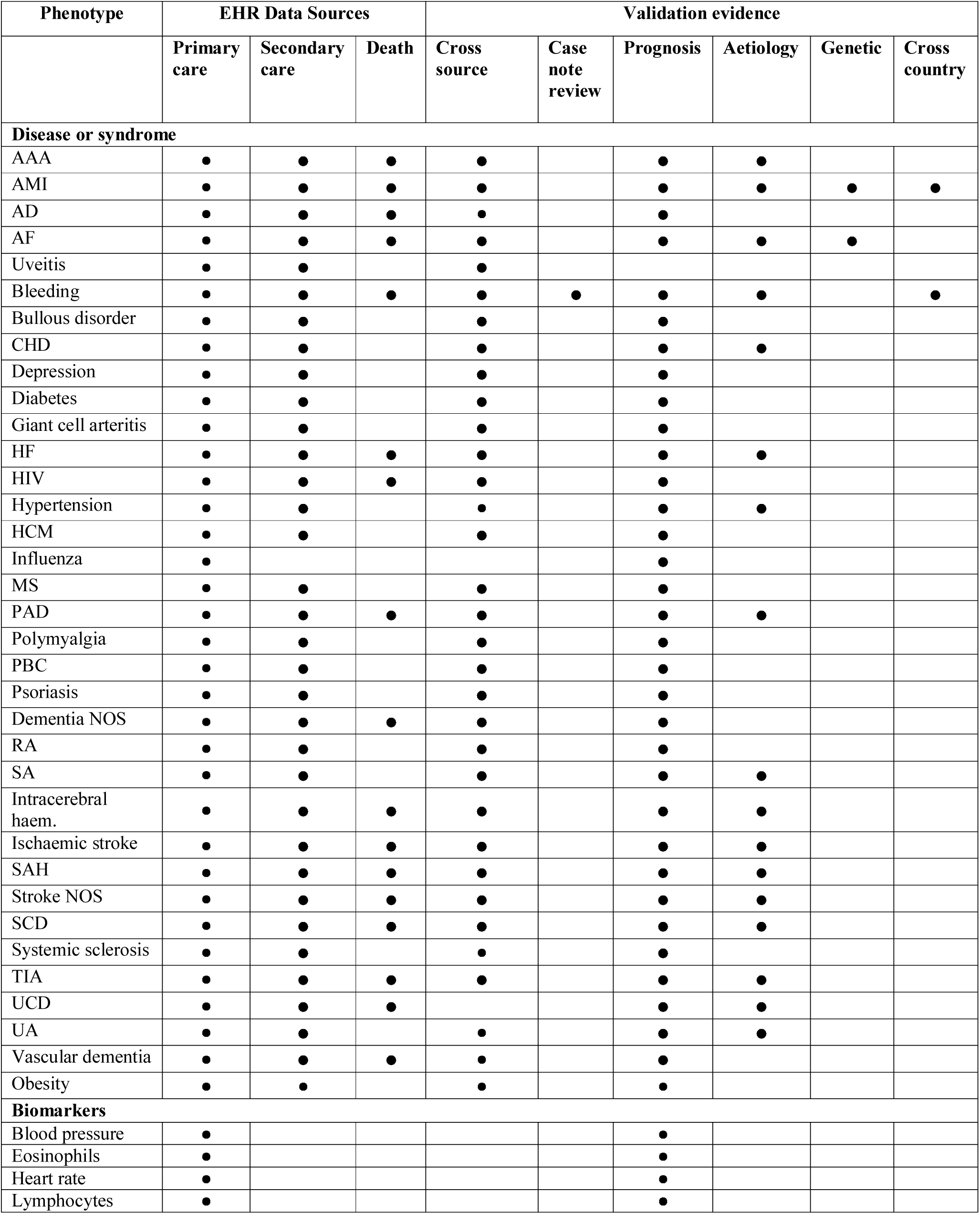
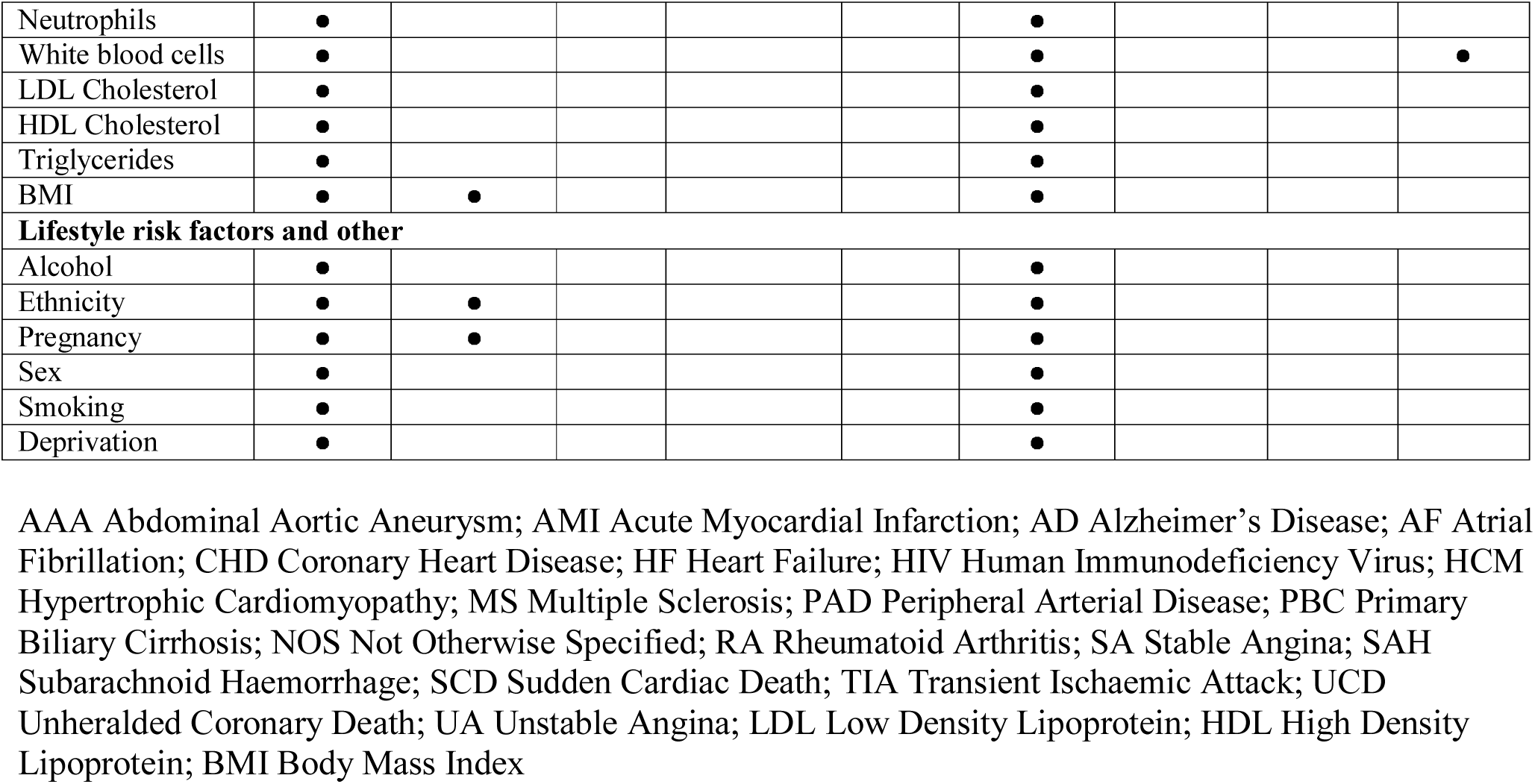
Overview of published, peer-reviewed EHR phenotypes derived from the CALIBER platform and the approaches of validation evidence - More information available on the CALIBER Portal https://www.caliberresearch.org/portal/phenotypes.

| Phenotype | EHR Data Sources |  |  | Validation evidence |  |  |  |  |  |
| --- | --- | --- | --- | --- | --- | --- | --- | --- | --- |
|  | Primary care | Secondary care | Death | Cross source | Case note review | Prognosis | Aetiology | Genetic | Cross country |
| **Disease or syndrome** |  |  |  |  |  |  |  |  |  |
| AAA | • | • | • | • |  | • | • |  |  |
| AMI | • | • | • | • |  | • | • | • | • |
| AD | • | • | • | • |  | • |  |  |  |
| AF | • | • | • | • |  | • | • | • |  |
| Uveitis | • | • |  | • |  |  |  |  |  |
| Bleeding | • | • | • | • | • | • | • |  | • |
| Bullous disorder | • | • |  | • |  | • |  |  |  |
| CHD | • | • |  | • |  | • | • |  |  |
| Depression | • | • |  | • |  | • |  |  |  |
| Diabetes | • | • |  | • |  | • |  |  |  |
| Giant cell arteritis | • | • |  | • |  | • |  |  |  |
| HF | • | • | • | • |  | • | • |  |  |
| HIV | • | • | • | • |  | • |  |  |  |
| Hypertension | • | • |  | • |  | • | • |  |  |
| HCM | • | • |  | • |  | • |  |  |  |
| Influenza | • |  |  |  |  | • |  |  |  |
| MS | • | • |  | • |  | • |  |  |  |
| PAD | • | • | • | • |  | • | • |  |  |
| Polymyalgia | • | • |  | • |  | • |  |  |  |
| PBC | • | • |  | • |  | • |  |  |  |
| Psoriasis | • | • |  | • |  | • |  |  |  |
| Dementia NOS | • | • | • | • |  | • |  |  |  |
| RA | • | • |  | • |  | • |  |  |  |
| SA | • | • |  | • |  | • | • |  |  |
| Intracerebral haem. | • | • | • | • |  | • | • |  |  |
| Ischaemic stroke | • | • | • | • |  | • | • |  |  |
| SAH | • | • | • | • |  | • | • |  |  |
| Stroke NOS | • | • | • | • |  | • | • |  |  |
| SCD | • | • | • | • |  | • | • |  |  |
| Systemic sclerosis | • | • |  | • |  | • |  |  |  |
| TIA | • | • | • | • |  | • | • |  |  |
| UCD | • | • | • |  |  | • | • |  |  |
| UA | • | • |  | • |  | • | • |  |  |
| Vascular dementia | • | • | • | • |  | • |  |  |  |
| Obesity | • | • |  | • |  | • |  |  |  |
| **Biomarkers** |  |  |  |  |  |  |  |  |  |
| Blood pressure | • |  |  |  |  | • |  |  |  |
| Eosinophils | • |  |  |  |  | • |  |  |  |
| Heart rate | • |  |  |  |  | • |  |  |  |
| Lymphocytes | • |  |  |  |  | • |  |  |  |
| Neutrophils | ● |  |  |  |  | ● |  |  |  |
| White blood cells | ● |  |  |  |  | ● |  |  | ● |
| LDL Cholesterol | ● |  |  |  |  | ● |  |  |  |
| HDL Cholesterol | ● |  |  |  |  | ● |  |  |  |
| Triglycerides | ● |  |  |  |  | ● |  |  |  |
| BMI | ● | ● |  |  |  | ● |  |  |  |
| **Lifestyle risk factors and other** |  |  |  |  |  |  |  |  |  |
| Alcohol | ● |  |  |  |  | ● |  |  |  |
| Ethnicity | ● | ● |  |  |  | ● |  |  |  |
| Pregnancy | ● | ● |  |  |  | ● |  |  |  |
| Sex | ● |  |  |  |  | ● |  |  |  |
| Smoking | ● |  |  |  |  | ● |  |  |  |
| Deprivation | ● |  |  |  |  | ● |  |  |  |

AAA Abdominal Aortic Aneurysm; AMI Acute Myocardial Infarction; AD Alzheimer's Disease; AF Atrial Fibrillation; CHD Coronary Heart Disease; HF Heart Failure; HIV Human Immunodeficiency Virus; HCM Hypertrophic Cardiomyopathy; MS Multiple Sclerosis; PAD Peripheral Arterial Disease; PBC Primary Biliary Cirrhosis; NOS Not Otherwise Specified; RA Rheumatoid Arthritis; SA Stable Angina; SAH Subarachnoid Haemorrhage; SCD Sudden Cardiac Death; TIA Transient Ischaemic Attack; UCD Unheralded Coronary Death; UA Unstable Angina; LDL Low Density Lipoprotein; HDL High Density Lipoprotein; BMI Body Mass Index

### 1) Cross-EHR source concordance

The PPV of AMI (the probability that the diagnosis recorded in MINAP was AMI rather than unstable angina or a non-cardiac diagnosis) was 92.2% (6,660/7,224, 95% CI 91.6%-92.8%) in CPRD and 91.5% (6,851/7,489, 90.8%-92.1%) in HES **(Figure 2)**. Among the 17,964 patients with at least one record of non-fatal AMI, 13,380 (74.5%) were recorded by CPRD, 12,189 (67.9%) by HES, and 9,438 (52.5%) by MINAP. Overall, 5,561 (31.0%) of patients had AMI recorded in three sources (32.0% within 90 days) and 11,482 (63.9%) in at least two sources. For 89,554 HF cases, 26% were recorded in CPRD only, 27% in both CPRD and HES, 34% in HES only, and 13% had HF as cause of death without a previous record elsewhere. In 39,804 bleeding cases 59.4% were captured in CPRD, 50.2% in HES, and 3.8% events in ONS. Allowing a 30-day window, only 13.2% of events were captured in two or more sources. Similarly, a very small proportion (13.2%) of bleeding cases identified were captured in multiple data sources.

### 2) Case note review

We tested the validity of ICD-10 terms used in our bleeding phenotype and found an NPV of 0.94 (0.90, 0.97) and a PPV of 0.88, i.e. 88% of bleeding events identified by the ICD-10 terms utilized in the CALIBER bleeding phenotype were indeed bleeding events according to the independent review of the entire hospital record by two clinicians, blinded to the term assignment. We found that ICD-10 coded events underestimate the occurrence of bleeding, with a sensitivity estimate of 0.48, consistent with a previous study where 38% of hospitalised bleeding events were not captured by coded terms [63]. Specificity was found to be 0.99 (0.97, 1.00) indicating a very low number of false positive bleeding events.

### 3) Aetiology

**Figure 3** shows age and sex adjusted HRs from Cox proportional hazards models for HF and CVD risk factors (smoking, Type-II diabetes, systolic blood pressure, heart rate) in CALIBER and non-EHR studies.

### 4) Prognosis

In 20,819 AMI cases we found that patients with events recorded in only one source had higher mortality than those recorded in more than one source (age and sex adjusted HR 2.29, 95% CI 2.17 to 2.42; P<0.001) [29]. Among patients with AMI recorded in only one source, those only in CPRD had the highest mortality on the first day but the lowest mortality thereafter. Among patients with AMI recorded in HES or MINAP, those in MINAP had lower coronary mortality in the first month (age and sex adjusted HR 0.33, 0.28 to 0.39, P<0.001) but similar mortality for non-coronary events (1.12, 0.90 to 1.40, P=0.3). After the first month, patients with AMI in CPRD had about half the hazard of mortality of patients with AMI recorded in one of MINAP or HES (age/sex adjusted HR 0.49, 95% CI 0.40-0.60, P<0.001). In 89,994 HF cases, we observed 51,903 deaths and generated KM curves for 90◻ day survival. Adjusted for age and sex, HF was strongly associated with mortality, with HRs for all◻ cause mortality ranging from 7.01 (95% CI 6.83–7.20), 7.23 (95% CI 7.03-7.43), up to 15.38 (95% CI 15.02-15.83) for patients in CPRD with acute HF hospitalization, CPRD only, and HES only, compared with a age/sex-matched reference population. Age, concomitant COPD, and diabetes were amongst the strongest predictors of death. Compared to patients with no bleeding, patients with bleeding recorded in CPRD and HES were at increased risk of all-cause mortality and atherothrombotic events. (HR all-cause mortality 1.98 (95% CI: 1.86, 2.11) for CPRD bleeding, and 1.99 (95% CI: 1.92, 2.05) for HES bleeding).

### 5) Genetic associations

In the CARDIoGRAMplusC4D GWAS summary data, we identified 31 independent variants associated with AMI by linkage disequilibrium (LD) clumping (R^2^<0.001, 250 kb) genetic variants reaching genome-wide significance (P <5×10−8). In the UK Biobank, we identified 8,281 AMI cases, 394,933 controls and excluded 5,266 participants from the analysis due to self-reported AMI at baseline. From 31 previously-reported SNPs, 31 (100%) had P<0.05 with same direction, with 26 (83.8%) passing Bonferroni correction (P<0.0016)(Supplementary Table 1).

### 6) External populations

We assessed the validity of the AMI, HF and bleeding phenotypes by comparing long-term outcomes (any cause death, fatal AMI/stroke, hospital bleeding) in AMI survivors in England (n=4,653), Sweden (n=5,484), US (n=53,909) and France (n=961) [64]. We found consistent associations with 12 baseline prognostic factors (age, gender, AMI, HF, diabetes, stroke, renal disease, peripheral arterial disease, atrial fibrillation, hospital bleeding, cancer, Chronic Obstructive Pulmonary Disease (COPD)) and each outcome. In each country, we observed high 3-year crude cumulative risks of all-cause death (from 19.6% [England] to 30.2% [US]); the composite of AMI, stroke, or death [from 26.0% (France) to 36.2% (US)]; and hospitalized bleeding [from 3.1% (France) to 5.3% (US)]. Adjusted risks were similar across countries, but higher in the US for all-cause death [RR US vs. Sweden, 1.14 (95% CI 1.04–1.26)] and hospitalized bleeding [RR US vs. Sweden, 1.54 (1.21–1.96)]. Similar analyses were performed for white blood cell (WBC) comparing all-cause mortality in England and New Zealand (NZ) [65,66]. High WBC within the reference range (8.65–10.05×10^9^/L) was associated with significantly increased mortality compared to the middle quintile (6.25–7.25×10^9^/L); adjusted HR 1.51 (95% CI 1.43-1.59) [England], 1.33 (95% CI 1.06-1.65) [NZ].

## DISCUSSION

In this study, we describe the CALIBER phenotyping approach which has been used to produce 51 validated phenotyping algorithms which capture disease status, biomarker values and lifestyle risk factors across four UK national EHR data sources spanning primary care, hospital admissions, a disease registry and a mortality register. Creating nationally-applicable phenotypes leveraging multiple EHR sources has, until recently, been a challenging, time-consuming, unreplicable and somewhat opaque process without any centralised resources. Research studies require precise phenotype definitions but phenotypic information found in EHR is typically inconsistent and of variable data quality. These problems are exacerbated when linking data across healthcare settings (i.e. primary care and hospital admissions) as each source records information using different healthcare processes, disparate formats and terminologies and interact with fundamentally different patient populations. Complementary initiatives exist [19] but these are different from CALIBER as they focus on curating codelists. We have taken a different approach and define an EHR phenotype as a combination of three essential elements: controlled clinical terminology terms, implementation logic and validation evidence all of which are curated on the CALIBER Portal. Compared with the Phenotype Knowledgebase (PheKB) developed by the eMERGE consortium, CALIBER includes additional approaches of validation e.g. aetiological and prognostic across population samples but lacks comprehensive detailed PPV/NPV measurements which are made possible by the availability of text and access to case notes at scale in the US.

CALIBER phenotyping algorithms use structured information on diagnoses, symptoms, referrals, prescriptions and procedures which are recorded using five controlled clinical terminologies and continuous measurements in order to extract disease onset and severity. The actual phenotyping algorithm production was lengthy and labor intensive and usually involved a large number of iterations although the exact number of person hours was difficult to ascertain. A particular challenge was the need to reconcile differences in granularity of diagnosis terms used in primary care and secondary care EHR as each healthcare setting utilizes different clinical terminologies to record information (Read in primary care, ICD-10 in secondary care). For example, in HF, the Read controlled clinical terminology allowed us to potentially distinguish between the two main congestive heart failure types: heart failure with normal/preserved ejection fraction (HFpEF) (i.e. Read term “G583.11 HFNEF - heart failure with normal ejection fraction”) and heart failure with reduced ejection function (HFrEF) / left ventricular systolic dysfunction (i.e. Read term “G5yy900 - Left ventricular systolic dysfunction”). Conversely, ICD-10 terms are substantially less specific (i.e. ICD terms “I50.0 Congestive heart failure” and “I50.9 Heart failure, unspecified”) and do not allow for this distinction. As a rule, for overlapping diagnoses across multiple sources, CALIBER phenotypes utilize the source with the highest clinical resolution to ascertain disease status.

We found that diagnosis terms in primary care using Read terms were not always informative and could not directly be used to ascertain particular phenotypes. For example, when attempting to create a dietary phenotype, we identified 173 Read terms related to nutrition which were recorded across 5.6M diagnosis events. Only 8% of these however were sufficiently informative to infer a particular nutritional diet i.e low fat diet, gluten free diet, diabetic diet or low sodium diet. The majority of terms used were generic terms which carried little information (i.e. “8CA4.00 Patient advised re diet” or “9N0H.00 Seen in dietician clinic”) and could not be used for ascertaining a phenotype with reasonable performance. While such terms could potentially be utilized as supporting information for other phenotypes (i.e. diabetic diet as evidence of diabetes) they cannot be used to ascertain a phenotype directly.

We observed that clinically-informed combinations of information across EHR sources improves case detection. All disease and syndrome phenotypes (n=35) utilized information sourced from primary care and hospital care EHR and roughly half (n=18) utilizing cause-specific mortality information recorded in the national death register. In general, we considered EHR sources complementary to each other with each providing a different aspect of a patient’s disease state (chronic vs. acute) rather than denote one as the authoritative source of information. One notable exception to this is mortality where we used the ONS date of death as the “gold standard” as studies have shown that discrepancies exist between the recorded death dates in primary care EHR and the date recorded on the death certificate (ONS). A previous study [67] of 118,571 deaths in England showed that a discrepancy existed in almost a quarter of deaths. Considerable variation was observed between GP practices on the degree of such discrepancies (in the majority of cases, the date of death recorded by the GP was after the date of death recorded on the death certificate). This is because GPs do not routinely see the death certificate (which is the definitive record of time and cause of death) and there is no legal obligation for them to record the date of death accurately. If there is a delay in their receipt of notification of death, they might record the date of death as the date of notification, or the date the patient’s record was closed, rather than the actual date of death. In line with previous literature we therefore used the ONS as the “gold standard” for ascertaining mortality.

A major effort of CALIBER has been to create longitudinal disease phenotypes that capture early and late manifestations of disease. We observed that the proportion of cases contributed by each EHR source differed by age at diagnosis: patients identified in secondary care EHR were substantially older than those identified in primary care. For example, substantially more, atrial fibrillation cases were identified in secondary care EHR rather than in primary care (32,930 cases compared to 11,068 from primary care) and using only a single source would have introduced bias and underestimated the incidence of disease. Conversely type 2 diabetes cases were exclusively identified through primary care EHR with no cases identified exclusively in hospital EHR due to the fact that, like other diseases such as hypertension, diagnosis and management is almost entirely performed in primary care.

Validation (Table 2) was a critical step for assessing the accuracy of EHR-derived phenotypes. We did not consider algorithm validation as a finite task but as a constantly evolving process due to the underlying complexity of EHR data and the processes which generate them [68]. We sought to address the spectrum of validation views and developed an approach which captures six levels of evidence. The majority of disease and syndrome phenotypes examined incidence estimates across different EHR sources and consistency with previous associations in terms of disease aetiology and prognosis. Validation was more restricted in biomarker and lifestyle risk factor phenotypes since information was derived from only one particular source (in the case of biomarkers, measurements were exclusively identified in primary care). Clinician case note review was considered the “gold standard” of phenotype validation that enables PPV/NPV calculations but access to medical records was not widely available and thus we could only perform this in a single instance. Prognostic validation was one of the main validation approaches where consistency with previously-reported findings provided a degree of confidence in terms of phenotype validity (for exposures, outcomes and covariates used in the analyses). Inconsistent results however were possible and could be explained due to multiple factors such as different health settings and sources of data, healthcare process factors and workflows and incomparable definitions.

**Table 2:** Systematic validation of the CALIBER EHR-derived phenotypes for a) heart failure, b) acute myocardial infarction, and c) bleeding across six approaches of evidence: cross-EHR concordance, case-note review, aetiology, prognosis, genetic associations and external populations.

| Validation domain | Description | What has been done |  |  |
| --- | --- | --- | --- | --- |
|  |  | Heart failure | Acute Myocardial Infarction | Bleeding |
| **Cross EHR source concordance** | To what extent is the phenotype concordant across EHR sources? | The proportion of HF cases recorded in primary care and hospital care EHR was 27% [31]. | The proportion of non-fatal AMI defined across primary care, hospital care and disease registry was 32% [29]. | The proportion of bleeding events recorded in primary care and hospital care was 12% with 47% of bleeding events recorded only in primary care and 12% only in hospital. |
| **Case-note review** | What is the positive predictive value and the negative predictive value when comparing the algorithm with clinician review of case notes or “gold standard” source of information? |  | Compared with AMI defined in the disease registry, the PPV of AMI recorded in primary care was 92.2% (95% CI 91.6% to 92.8%) and in hospital admissions was 91.5% (90.8% to 92.1%) [29] | Compared through independent review by two clinicians, the PPV of bleeding events identified through the phenotyping algorithm was 0.88. |
| **Aetiology** | Are the prospective associations with risk actors consistent with previous evidence? | Type 2 diabetes [84], systolic/diastolic blood pressure [32], heart rate [85], socioeconomic deprivation [86], alcohol consumption [87], smoking [88], ethnicity [44], acute myocardial infarction [29], depression [89], neutrophil counts [90], eosinophil/lymphocyte counts [91], atrial fibrillation [30], sex [92] | Type 2 diabetes [84], blood pressure [32], heart rate [85], socioeconomic deprivation [86], alcohol consumption [87], smoking [88], ethnicity [44], acute myocardial infarction [29], depression [89], neutrophils [90], eosinophil/lymphocyte counts [91], atrial fibrillation [30], influenza infection [93], ischaemic presentations [94], sex [92] | At 5 years 29.1% (95% CI: 28.2, 29.9%) of atrial fibrillation patients, 21.9% (21.2, 22.5%) of myocardial infarction patients, 25.3% (24.2, 26.3%) of unstable angina patients and 23.4% (23.0, 23.8%) of stable angina had bleeding of any kind |
| **Prognosis** | Are the risks of subsequent events plausible? | Corrected for age and sex, HF was strongly | Patients with myocardial infarction identified | The hazard ratio for all-cause mortality was 1.98 (95% CI: |
|  |  | associated with mortality, with HRs for all-cause mortality ranging from 7.01 (95% CI 6.83–7.20), 7.23 (95% CI 7.03–7.43), up to 15.38 (95% CI 15.02–15.83) for patients in primary care with acute HF hospitalization, primary care only, and patients hospitalized but no PC record [31]. | in the disease registry had lower crude 30-day mortality (10.8%, 95% CI 10.2% to 11.4%) than those identified in hospital care (13.9%, 13.3% to 14.4%) or in primary care (14.9%, 14.4% to 15.5%, fig 2). At one year, however, mortality was similar in all three groups, at around 20% [29]. Of the 24,479 patients with AMI, 5775 (23.6%) developed HF during a median follow-up of 3.7 years (incidence rate per 1000 person-years: 63.8, 95% CI 62.2 to 65.5) [95] | 1.86, 2.11) for primary care bleeding with markers of severity, and 1.99 (95% CI: 1.92, 2.05) for hospitalised bleeding without markers of severity, compared to patients with no bleeding. |
| **Genetic associations** | Are the observed genetic associations plausible and concordance with previous evidence? |  | Consistent direction and magnitude of associations were replicated in 67 (97%) of previously reported genetic variants [4]. |  |
| **External populations** | Has the algorithm been tested (in any of the above validation domains) in different countries? |  | We observed high 3-year crude cumulative risks of all-cause death (from 19.6% [England] to 30.2% [USA]); the composite of AMI, stroke, or death [from 26.0% (France) to 36.2% (USA)]; and hospitalized bleeding [from 3.1% (France) to 5.3% (USA)]. [64] |  |

In terms of the complete hospital interaction, HES data are a snapshot of the patient journey as data are collected for hospital reimbursement [8,52]. Hospital records are converted into diagnosis and procedure codes locally (following an existing protocol) at each hospital and submitted to the NHS. HES data are provided to researchers with identifiers for hospitals removed in order to protect patient anonymity as a common identifier is used across HES and CPRD GP practices which have a substantially smaller catchment area. As such, we were unable to assess the effect of site-level variability in terms of data capture and quality and phenotype validity. Multiple initiatives however exist for obtaining raw hospital records for research e.g. National Institute for Health Research Health Informatics Collaborative (NIHR HIC) which links eleven Intensive Care Units (ICUs) in five hospitals for research (>18,000 patients, >21,000 admissions, median 1,030 time-varying measures [69]). Crucially, these initiatives will enable researchers to have access to raw hospital data, including free text, for creating and validating phenotypes and will create a feedback loop with clinical care that will provide detailed information on the healthcare processes generating the data (critical for phenotyping) and drive data standardization and quality.

CALIBER phenotyping algorithms are rule-based, deterministic, and provide a framework on which future phenotypes can be created. While our approach yields robust and accurate algorithms, it does not scale with our ambition to create and curate thousands of high-quality phenotypes (and their validation) that capture the entire human phenome. To do this, research is required on high-throughput phenotyping involving supervised and unsupervised learning approaches and natural language processing[70]. Such methods can generate thousands of phenotypes and discover hidden associations within clinical data in a fraction of the current cost and time requirements and with minimal human intervention. Robust approaches for classifying phenotype complexity are required to ensure proportional resourcing for phenotyping [71]. Finally, a key next step is to use the six sites of the recently funded Health Data Research UK (HDR UK) national institute in order to scale up the number of phenotypes created and curated using UK EHR.

The use of a common data model to map between the clinical terminologies used across EHR sources, such as the Observational Medical Outcomes Partnership (OMOP) Common Data Model (CDM) can potentially address some of the labour-intensive tasks associated with phenotyping. For example, the translation from phenotype definition to SQL for data extraction was manual due to the lack of an established storage format [72] for the algorithms and variable schema across EHR sources. OMOP CDM can potentially act as Relational Database Management System (RDBMS) agnostic schema which standardized analytical tools can be deployed on and has been shown to be robust [73,74] and we are currently in the process of evaluating the fidelity of the data transformation. We have additionally evaluated different approaches (Semantic Web Technologies, openEHR [75,76]) for storing phenotype definitions in a computable format that can enable high-throughput phenotyping and eliminate the need for manual human-driven translation to SQL queries. Given that all of UK primary care EHR data are hosted on four clinical information systems vendors, there is a real opportunity to create computable phenotypes which can be utilized across the NHS [77]. To accomplish this, information exchange standards (e.g. Fast Healthcare Interoperability Resources (FHIR) [78]) have to be utilized and combined with approaches such as the Phenotype Execution and Modeling Architecture (PHEMA)[79] and the Measure Authoring Tool (MAT) [80].

## CONCLUSION

We have demonstrated the strengths and challenges of phenotyping national UK EHR data using three exemplars (HF, AMI, bleeding) and have exemplified the UK’s national EHR phenomics platform. In this manuscript, we presented the CALIBER platform as a framework for using national EHR from primary and secondary health care, disease and national mortality registries. CALIBER is analogous and complementary to other leading initiatives, e.g. eMERGE, in that it ensures best practice and reproducibility for creating and validating EHR-derived phenotypes [81,82]. In contrast with eMERGE however, which uses secondary care data (higher proportion of disease), CALIBER exploits primary care EHR which contain healthy and ill individuals. Importantly, the approaches described here are potentially scalable/adaptable to the entire 65M UK population and is work in underway to create a chronological map of human disease spanning early and late life by curating over 300 diseases [96].

Through CALIBER we provide a framework for the consistent definition, use and re-use of EHR derived phenotypes from national UK EHR sources for observational research: a) high-resolution clinical epidemiology using national EHR examining disease aetiology or prognosis, or b) genetic epidemiology studies through the UK Biobank and Genomics England investigating simple and complex traits across populations. One of the primary audiences of CALIBER phenotypes is international: US investigators account for a third of studies using UK primary care EHR [18] and two thirds of UK Biobank studies are carried out by US investigators. Additionally, the controlled clinical terminologies used in UK EHR are directly translatable and transferable to the US e.g. Read terms (CTV3 (Clinical Terms Version 3) are part of SNOMED-CT, and ICD-9-Clinical Modification (CM) to ICD-10 mappings exist. As PheKB and other initiatives evolve, establishing links across national portals [83] and infrastructure can enable cross-biobank analyses of complex/rare phenotypes [7].

The creation of a national phenomics platform through CALIBER provides an opportunity for the UK EHR community to interact, nationally and internationally, and connects data producers and consumers. Researchers can deposit phenotyping algorithms in the Portal for others to download, refine and use. EHR users i.e. clinicians can view these algorithms and understand how the data they record is being used for research.

## ACKNOWLEDGEMENTS

This study was approved by the Medicines and Healthcare Products Regulatory Agency (MHRA) Independent Scientific Advisory Committee (ISAC) - protocol references: 11_088, 12_153R, 16_221, 18_029R2, 18_159R.

This study is based in part on data from the Clinical Practice Research Datalink obtained under licence from the UK Medicines and Healthcare products Regulatory Agency. The data is provided by patients and collected by the NHS as part of their care and support. The interpretation and conclusions contained in this study are those of the author/s alone.

Hospital Episode Statistics Copyright © (2019), re-used with the permission of The Health & Social Care Information Centre. All rights reserved.

The OPCS Classification of Interventions and Procedures, codes, terms and text is Crown copyright (2016) published by Health and Social Care Information Centre, also known as NHS Digital and licensed under the Open Government Licence available at www.nationalarchives.gov.uk/doc/open-government-licence/open-government-licence.htm

This study was carried out as part of the CALIBER programme (https://www.ucl.ac.uk/health-informatics/caliber). CALIBER, led from the UCL Institute of Health Informatics, is a research resource consisting of linked electronic health records phenotypes, methods and tools, specialised infrastructure, and training and support.

The views expressed are those of the author(s) and not necessarily those of the NHS, the NIHR or the Department of Health.

