## Supplementary figures and images for "UK phenomics platform for developing and validating EHR phenotypes: CALIBER"

### Figure 1

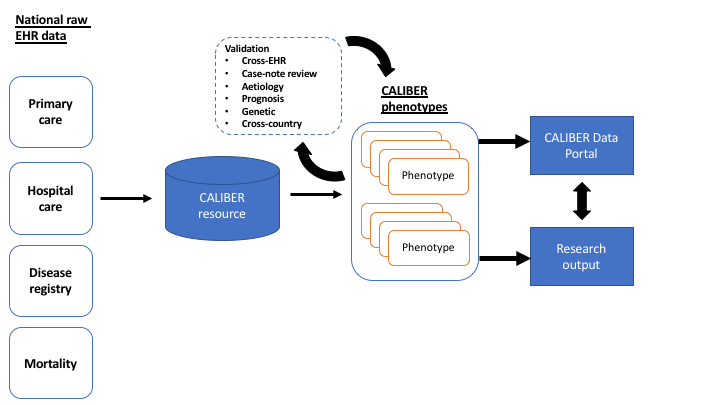

### Figure 2

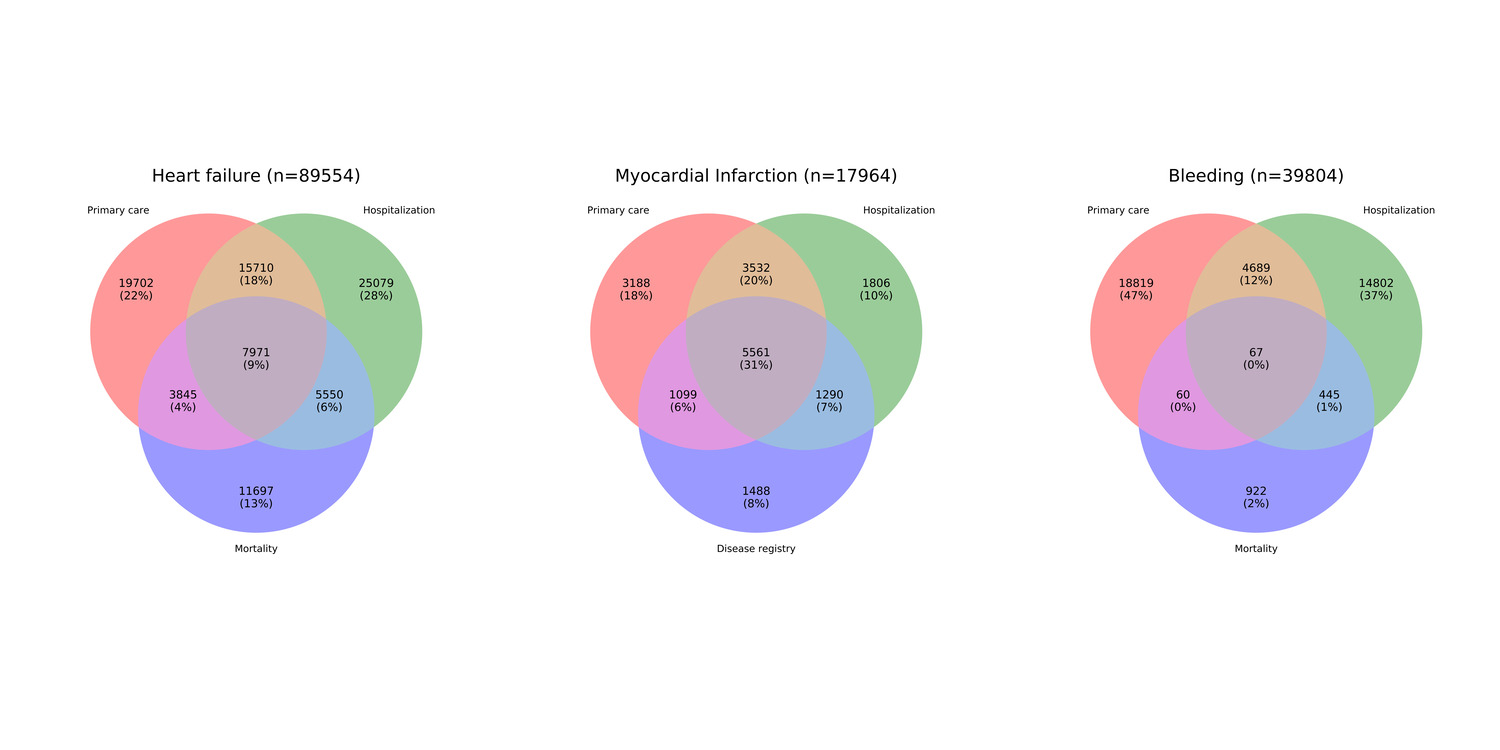

### Figure 3

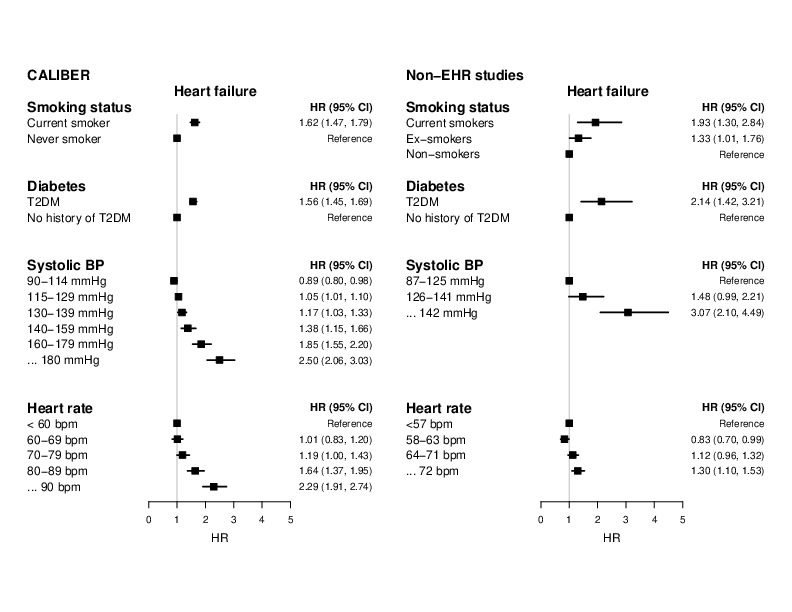

### Figure 4

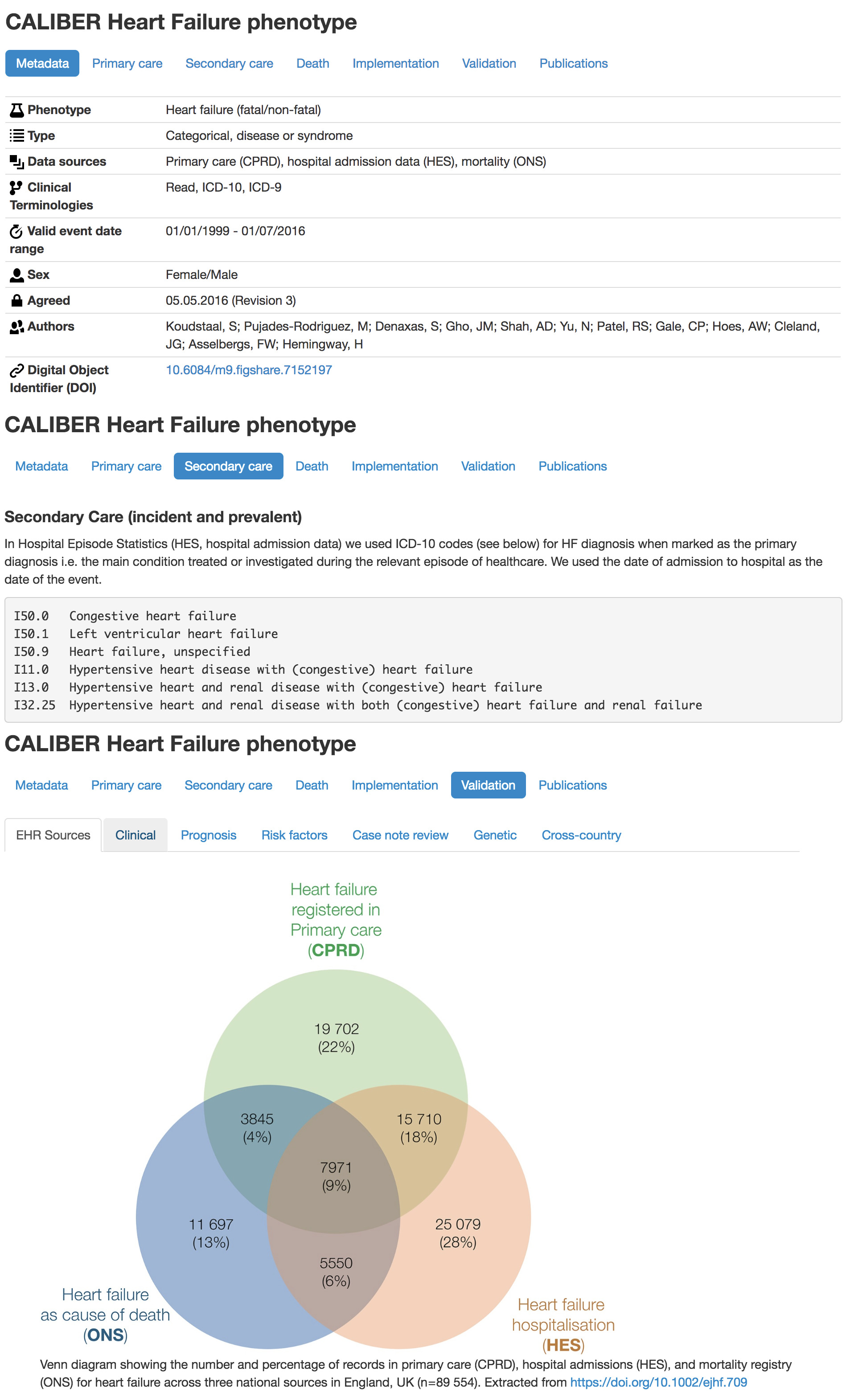
