## Supplemental Table 1 for "UK phenomics platform for developing and validating EHR phenotypes: CALIBER"

**Supplementary Table 1: Single Nucleotide Polymorphisms (SNPs) associated with acute myocardial infarction in CARDIoGRAMplusC4D and p-values obtained from replication experiment in the UK Biobank.**
Table is restricted to SNPs reaching genome-wide significance (P <5x10^-8^) in CARDIoGRAMplusC4D**.** SNP Single Nucleotide Polymorphism; CHR Chromosome; BP Base-pair coordinate; A1 Minor allele; A2 Major allele; C4D CARDIoGRAMplusC4D; SE Standard Error (of beta) estimate; UKB UK Biobank

| **SNP** | **CHR** | **BP** | **A1** | **A2** | **C4D beta** | **C4D SE** | **C4D p-value** | **UKB beta** | **UKB SE** | **UKB p-value** |
| --- | --- | --- | --- | --- | --- | --- | --- | --- | --- | --- |
| rs10176176 | 2 | 85762048 | T | A | 0.0646494 | 0.0102571301294879 | 2.92178112635262e-10 | 0.0704584636485614 | 0.01601 | 1.071e-05 |
| rs10455872 | 6 | 161010118 | G | A | 0.284774 | 0.0265918442181448 | 9.22949771139045e-27 | 0.315540400580177 | 0.02604 | 9.36e-34 |
| rs10947786 | 6 | 39156410 | A | G | -0.0718678 | 0.0127578871999083 | 1.76902575065445e-08 | -0.0766650849354854 | 0.0199 | 0.0001168 |
| rs113113862 | 19 | 11183577 | A | G | -0.075448 | 0.0126926448335797 | 2.77780375840091e-09 | -0.0542450148523958 | 0.01856 | 0.003449 |
| rs114155121 | 2 | 203718682 | C | T | 0.123752 | 0.0168087084645564 | 1.80619322659418e-13 | 0.144100343973757 | 0.02261 | 1.731e-10 |
| rs11556924 | 7 | 129663496 | T | C | -0.0689191 | 0.0125520732877071 | 4.00460798823928e-08 | -0.0609184149694171 | 0.01635 | 0.0001939 |
| rs1332329 | 10 | 91003419 | C | A | 0.0791447 | 0.0108278720670352 | 2.68490171929427e-13 | 0.0544881852840698 | 0.01654 | 0.000961 |
| rs143843429 | 6 | 161383079 | G | A | 0.325165 | 0.0458616278007066 | 1.33974463844549e-12 | 0.342170257735851 | 0.05894 | 6.369e-09 |
| rs17696736 | 12 | 112486818 | G | A | 0.0660518 | 0.0116368559482056 | 1.37819614625793e-08 | 0.113328685307003 | 0.01594 | 1.352e-12 |
| rs180803 | 22 | 24658858 | T | G | -0.186597 | 0.0316322462431263 | 3.65877089870357e-09 | -0.189104509090787 | 0.08307 | 0.02283 |
| rs1870634 | 10 | 44480811 | G | T | 0.0696961 | 0.0107321546480136 | 8.35200135321394e-11 | 0.0680647309141294 | 0.01702 | 6.35e-05 |
| rs2019090 | 11 | 103668962 | T | A | -0.0654617 | 0.0110961383759523 | 3.64656224882943e-09 | -0.0751074724868055 | 0.01729 | 1.403e-05 |
| rs2327426 | 6 | 134202690 | C | T | -0.0625017 | 0.0109700532107612 | 1.21587307318127e-08 | -0.0862116968190654 | 0.01765 | 1.036e-06 |
| rs2505083 | 10 | 30335122 | C | T | 0.0612049 | 0.0105660548801845 | 6.93072160959532e-09 | 0.0392207131532813 | 0.01597 | 0.01353 |
| rs2681472 | 12 | 90008959 | G | A | 0.0727045 | 0.0125053949499555 | 6.10530199183567e-09 | 0.0582689081239758 | 0.02074 | 0.004992 |
| rs28451064 | 21 | 35593827 | A | G | 0.122318 | 0.0177856270551088 | 6.09652233505703e-12 | 0.138891998866619 | 0.02336 | 2.96e-09 |
| rs35700460 | 1 | 222811407 | G | A | 0.0818302 | 0.0121908795293811 | 1.9140681993339e-11 | 0.110819835105244 | 0.01798 | 7.081e-10 |
| rs41290120 | 19 | 45382675 | A | G | -0.185714 | 0.0311929333356437 | 2.62138111683436e-09 | -0.127719741602371 | 0.0389 | 0.001031 |
| rs4773141 | 13 | 110954353 | G | C | 0.0802207 | 0.0128784997323974 | 4.69275570946371e-10 | 0.0610950993598108 | 0.01704 | 0.0003257 |
| rs4977574 | 9 | 22098574 | G | A | 0.188677 | 0.0102921335174754 | 4.58369089578329e-75 | 0.211070970079941 | 0.01592 | 3.553e-40 |
| rs532436 | 9 | 136149830 | A | G | 0.110862 | 0.0130821675141273 | 2.36483756955355e-17 | 0.0723206615796261 | 0.02009 | 0.0003292 |
| rs627135 | 10 | 44765000 | G | T | -0.0971445 | 0.0155173817599531 | 3.84099330268284e-10 | -0.141333176116644 | 0.02796 | 4.287e-07 |
| rs653178 | 12 | 112007756 | T | C | -0.0770565 | 0.0115792323450502 | 2.8386935060469e-11 | -0.12751332029896 | 0.01587 | 9.217e-16 |
| rs7173743 | 15 | 79141784 | C | T | -0.0639689 | 0.0104014466730329 | 7.74845818846077e-10 | -0.0643252105565576 | 0.016 | 5.829e-05 |
| rs72689147 | 4 | 156639888 | T | G | -0.0735497 | 0.0130276558001553 | 1.64569440110862e-08 | -0.05508996631796 | 0.02076 | 0.007924 |
| rs72934535 | 2 | 203968973 | C | T | 0.141086 | 0.0186043758299238 | 3.36350520228249e-14 | 0.136277618292548 | 0.02476 | 3.674e-08 |
| rs7528419 | 1 | 109817192 | G | A | -0.101314 | 0.0126204275432192 | 9.92458954724651e-16 | -0.118558336124135 | 0.01963 | 1.57e-09 |
| rs9295128 | 6 | 160751531 | T | G | 0.484677 | 0.0523424052915257 | 2.04911126602025e-20 | 0.471252848646168 | 0.05222 | 1.897e-19 |
| rs9349379 | 6 | 12903957 | G | A | 0.130965 | 0.0106499256590043 | 9.36640553554371e-35 | 0.106160195828391 | 0.01602 | 3.637e-11 |
| rs9457761 | 6 | 160292603 | G | C | 0.317096 | 0.0568644095011557 | 2.45615612968301e-08 | 0.204572165728774 | 0.05783 | 0.0004142 |
| rs9970807 | 1 | 56965664 | T | C | -0.110641 | 0.0183978107720577 | 1.81204059386386e-09 | -0.102143473668704 | 0.02861 | 0.0003591 |
